# Discovery and characterization of H_v_1-type proton channels in reef-building corals

**DOI:** 10.1101/2021.04.08.439075

**Authors:** Gisela E. Rangel-Yescas, Cecilia Cervantes, Miguel A. Cervantes-Rocha, Esteban Suarez-Delgado, Anastazia T. Banaszak, Ernesto Maldonado, Ian. S. Ramsey, Tamara Rosenbaum, León D. Islas

**Affiliations:** Departmento de Fisiología, Facultad of Medicina, Universidad Nacional Autónoma de México, Mexico City, Mexico; Unidad Académica de Sistemas Arrecifales, Instituto de Ciencias del Mar y Limnología, Universidad Nacional Autónoma de México, Puerto Morelos, Quintana Roo, Mexico; EvoDevo Research Group, Unidad Académica de Sistemas Arrecifales, Instituto de Ciencias del Mar y Limnología, Universidad Nacional Autónoma de México, Puerto Morelos, Quintana Roo, Mexico; Department of Physiology and Biophysics, School of Medicine, Virginia Commonwealth University, Richmond, VA, USA; Departmento of Neurociencia Cognitiva, Instituto de Fisiología Celular, Universidad Nacional Autónoma de México, Mexico City, Mexico

## Abstract

Voltage-dependent proton-permeable channels are membrane proteins mediating a number of important physiological functions. Here we report the presence of a gene encoding for H_v_1 voltage-dependent, proton-permeable channels in two species of reef-building corals. We performed a characterization of their biophysical properties and found that these channels are fast-activating and modulated by the pH gradient in a distinct manner. The biophysical properties of these novel channels make them interesting model systems. We have also developed an allosteric gating model that provides mechanistic insight into the modulation of voltage-dependence by protons. This work also represents the first functional characterization of any ion channel in scleractinian corals. We discuss the implications of the presence of these channels in the membranes of coral cells in the calcification and pH regulation processes and possible consequences of ocean acidification related to the function of these channels.

## Introduction

Scleractinian or stony corals are organisms in the phylum Cnidaria that deposit calcium carbonate in the form of aragonite to build an exoskeleton. Stony corals are the main calcifying organisms responsible for the construction of coral reefs, which are major ecosystems hosting numerous and diverse organisms. Coral reefs also act as natural barriers from strong ocean currents, waves and tropical storms, providing coastal protection. This protection centers on the ability of scleractinian corals to produce enough calcium carbonate (CaCO_3_). The increase in atmospheric CO_2_ concentrations as a result of human activity poses threats to coral-reef building organisms due to rising sea surface temperatures (Hoegh-Guldberg, 1999) and because CO_2_ is taken up by the ocean, dangerously lowering the pH of the sea water (Caldeira and Wickett, 2003).

It is known that precipitation of the aragonitic form of calcium carbonate is facilitated at elevated pH values, at very low concentrations of protons. Calcification by scleractinian corals is a process that has been shown to be modulated by the pH of the solution in which calcium carbonate is precipitated (Allemand et al., 2011). To this end, corals produce a specialized compartment between the ectoderm and the external substrate or skeleton called calicoblastic compartment, which contains a fluid derived from the surrounding sea water. The composition of this calicoblastic fluid or liquor is strictly regulated by the coral to maintain both an elevated pH, often close to one unit higher than the surrounding sea water, and an increased concertation of Ca^2+^ and carbonates. The molecular details of pH regulation in the calicoblastic fluid are not understood completely. Involvement of proton-pumps has been postulated and is likely to be part of proton transport in corals. Both P-type and V-type hydrogen pumps are present in coral transcriptomes and are known to play roles in the physiology of coral-algal symbiosis (Tresguerres et al., 2017). V-type H^+^-ATPases have also been shown to be involved in calcification in foraminifera (Toyofuku et al., 2017). If a proton pump is involved in lowering proton concentration in the calicoblastic fluid to maintain high calcification rates, protons will be transported to the cytoplasm of the ectodermal cells that constitute the calicoblastic epithelium, producing a profound acidification of the cytoplasmic pH (pH_i_). Although measurements of the (pH_i_) in corals indicate values of 7.13-7.4 (Venn et al., 2009), it is unknown how coral cells regulate pH_i_. Thus, an efficient pH-regulatory mechanism is to be expected to be present in corals. We hypothesized that proton channels might be fundamental to this physiological process and also required for calcification in hard corals.

Although a number of studies have delineated the physiological roles of H_v_1 voltage-gated proton channels in vertebrate cells (DeCoursey, 2013), less is known about their role in invertebrates. These channels are potential mediators in processes that are critically dependent on proton homeostasis. As an example, they have been shown to be involved in regulating the synthesis of the calcium carbonate skeleton in coccolithophores, calcifying unicellular phytoplankton (Taylor et al., 2011).

The range of voltages over which channel activation occurs is strongly modulated by the transmembrane proton gradient, characterized by: ΔpH = pH_o_-pH_i_ i.e., the difference between external and internal pH. In the majority of known H_v_1 channels, the voltage at which half the channels are activated, the V_0.5_ or the apparent threshold for channel opening (V_Thr_), shifts by roughly 40 mV per unit of ΔpH. Thus, the pH gradient strongly biases the voltage-independent free energy of channel activation (Cherny et al., 1995). With few exceptions, channel activation occurs at voltages that are more positive than the reversal potential for protons, implying that protons are always flowing outward under steady-state conditions. The fact that most H_v_1s mediate outward currents is the reason these channels are mostly involved in reversing intracellular acidification or producing voltage-dependent cytoplasmic alkalization (Lishko and Kirichok, 2010; DeCoursey, 2013).

Here we report the presence of genes encoding for H_v_1 channels in two species of reef-building corals. We cloned and characterized the biophysical properties of these channels in an expression system using patch-clamp electrophysiology. The demonstration of the presence of voltage-gated proton channels in corals is an initial step to a deeper understanding of coral calcification and its dysregulation under ocean acidification conditions. We show that some of the coral H_v_1’s biophysical properties are different from other known proton channels and this behavior makes them interesting models to try to understand some basic biophysical mechanisms in these channels. To explain this behavior, we develop a novel activation model to describe voltage- and pH-dependent gating that has general applicability to H_v_1 channels.

## Materials and methods

### Identification of H_v_1 sequences and cloning

Blast searches of the transcriptome of the Indo-Pacific coral *Acropora millepora* (Moya et al., 2012) detected four sequences that we identified as belonging to a putative proton-permeable channel. The GenBank accession numbers for these are: XM_015907823.1, XM_015907824.1, XM_029346499.1 and XM_029346498.1. We designed two pairs of oligonucleotides to amplify two of these sequences (Table 1). Total RNA was extracted from tissue obtained from a fragment of *Acropora millepora* acquired from a local salt-water aquarium provider. RNA was extracted by dipping the whole fragment for 2 min in 5 ml of solution D (4 M guanidinium thiocyanate, 25 mM sodium citrate, 5 % sarkosyl and 0.1 M 2-mercaptoethanol). After incubation, tissue was removed by gently pipetting the solution for 2 min. At this point, the calcareous skeleton was removed and RNA extraction continued according to (Chomczynski and Sacchi, 1987). Total RNA (1 µg) from *A. millepora* was used for RT-PCR, employing oligo dT and SuperScripII reverse transcriptase (Invitrogen, USA). Complementary DNA obtained from RT-PCR was used in three PCR reactions using oligos: 1) AcHv1Nter5’
s and 3’; 2) AcHv1Cter5’ and 3’ and, 3) AcHv1Nter5’ and AcHv1Cter3’ (Table 1). The Platinium Pfx DNA polymerase (Invitrogen) was used for amplification according to the manufacturer’s instructions. 1 µl of Taq DNA polymerase (Invitrogen, USA) was used for 10 min at 72 °C to add a poly A tail at 5’ and 3’ ends and facilitate cloning into the pGEM-T vector.

**Table 1.**
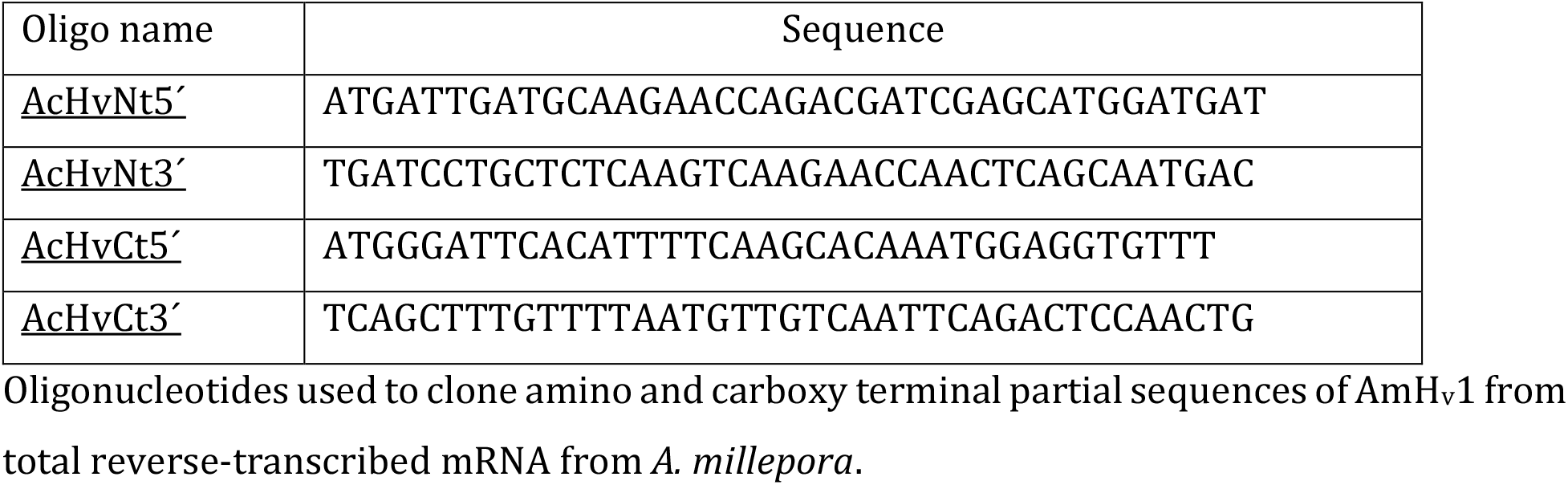

The PCR reaction 3 gave rise to a full open reading frame (ORF) containing AmH_v_1. New oligos AcHv1Nter5’
s and AcHv1Cter3’ containing restriction sites Kpn1 and Not1 respectively were used to re-amplify the ORF in pGEM-T and subclone it into pcDNA3.1 for heterologous expression.

The H_v_1 channel from *A. palmata* was cloned from a fragment of an adult specimen collected in the Limones Reef off of Puerto Morelos, Mexico. RNA extraction from small coral pieces was carried out by flash freezing in liquid nitrogen and grinding the frozen tissue. All other cloning procedures were as for *A. millepora*. All clones were confirmed by automatic sequencing at the Molecular Biology Facility of the Instituto de Fisiología Celular at UNAM.

### Heterologous expression of AmH_v_1

The cloned AmH_v_1 was expressed in HEK293 cells. Cells were grown on 100 mm culture dishes with 10 ml of Dulbecco’
ss Modified Eagle Medium (DMEM, Invitrogen) containing 10 % fetal bovine serum (Invitrogen, USA) and 100 units/ml-100 μg/ml of penicillin-streptomycin (Invitrogen, USA), incubated at 37°C in an incubator with 5.2 % CO_2_ atmosphere. When cells reached 90 % confluence, the medium was removed, and the cells were treated with 1 ml of 0.05 % Trypsin-EDTA (Invitrogen, USA) for 5 min. Subsequently, 1 ml of DMEM with 10 % FBS was added. The cells were mechanically dislodged and reseeded in 35 mm culture dishes over 5×5 mm coverslips for electrophysiology or in 35 mm glass bottom dishes for FRET experiments. In both cases, 2 ml of complete medium were used. Cells at 70 % confluence were transfected with pcDNA3.1-AmH_v_1 prepared from a plasmid midiprep, using jetPEI transfection reagent (Polyplus Transfection, Frence). For patch-clamp experiments, pEGFP-N1 (BD Biosciences Clontech, USA) was cotransfected with the channel DNA to visualize successfully transfected cells via their green fluorescence. Electrophysiological recordings were done one or two days after transfection.

### FRET measurement of stoichiometry

In order to measure the stoichiometry of subunit interaction employing FRET, we constructed fusion proteins between AmH_v_1 and mCerulean and mCitrine fluorescent proteins (FPs), to be used as donor and acceptor, respectively. The FPs were fused to the N-terminus of the channel in order to disrupt as little as possible the C-terminus mediated interaction. These constructs were transfected into HEK293 cells as described above. The apparent FRET efficiency between FP-containing constructs, *E*_*app*_, was measured via sensitized emission of the acceptor, employing the spectral-FRET method (De-la-Rosa et al., 2013; Zheng et al., 2002). Fluorescence was measured in a home-modified TE-2000U inverted epifluorescense microscope (Nikon, Japan). The excitation light source was an Argon Ion laser (Spectra-Physics, Germany) mainly producing light at 458, 488 and 514 nm; the laser beam is focused and then collimated using a 3 mm ball lens and a 50 mm focal length planoconvex lens. Collimated light is steered with a mirror and then is focused into the objective back focal plane by a 300 mm focal length achromatic lens.

Cells were imaged with a Nikon 60X oil immersion objective (numerical aperture 1.4). The detection arm of the microscope is coupled to a spectrograph (Acton Instruments, USA) and an EM-CCD camera (Ixon, Andor, Ireland) controlled by Micromanager software (Edelstein et al., 2014). Excitation was achieved with appropriate excitation filters (Chroma, Vermont, USA) for mCerulean (458 nm) and mCitrine (488 nm). The emission filter was a long-pass filter in order to collect the full emission spectrum of the FRET pair. Apparent FRET efficiency is plotted as a function of the fluorescence intensity ratio (I_donor_/I_acceptor_). This relationship can be fitted with models of subunit association with fixed stoichiometry, according to (De-la-Rosa et al., 2013).

### Electrophysiology

All chemicals for solutions were acquired from Sigma-Aldrich (Mexico). Proton current recordings were made from HEK293 cells expressing pCDNA3.1-AmHv1 in the inside-out, whole-cell and outside-out configurations of the patch-clamp recording technique. For whole-cell and inside-out recordings, the extracellular solution (bath and pipette, respectively) was (in mM): 80 TMA-HMESO_3_, 100 buffer (MES: pH 5.5, 6.0 and 6.5; HEPPES: pH 7.0, 7.5), 2 CaCl_2_, 2 MgCl_2_ and pH adjusted NMDG/TMAOH and HCl. The intracellular solution (pipette and bath respectively) was (in mM): 80 TMA-HMESO_3_, 100 buffer (MES: pH 5.5, 6.0 and 6.5; HEPES: pH 7.0, 7.5), 1 EGTA and pH adjusted NMDG/TMAOH and HCl.

### Conditions for recording zinc effects

The effect of zinc was evaluated in outside-out patches at a ΔpH of 1. The bath solution composition was (in mM): 100 TMA-HMESO_3_, 100 HEPES, 8 HCl, 2 CaCl_2_, 2 MgCl_2_, and the indicated concentration of ZnCl_2_. The pipette solution was (in mM): 100 TMA-MESO_3_, 100 MES, 8 HCl, 10 EGTA and 2 MgCl_2_. Both solutions were adjusted to pH 7 and pH 6 respectively with TMA-OH/HCl. Patches were placed in front of a perfusion tube that was gravity-fed with the appropriate solution. Tubes were changed with a home-built rapid perfusion system.

Macroscopic currents were low-pass filtered at 2.5 kHz, sampled at 20 kHz with an Axopatch 200B amplifier (Axon Instruments, USA) using an Instrutech 1800 AD/DA board (HEKA Elektronik, Germany) or an EPC-10 amplifier (HEKA Elektronik, Germany). Acquisition control and initial analysis was done with PatchMaster software. Pipettes for recording were pulled from borosilicate glass capillaries (Sutter Instrument, USA) and fire-polished to a resistance of 4-7 MΩ when filled with recording solution for inside- and outside-out recordings and 1-3 MΩ for whole-cell. The bath (intracellular) solutions in inside-out patches were changed using a custom-built rapid solution changer. For whole-cell recordings all the bath solution was exchanged to manipulate pH. In some recordings, linear current components were subtracted using a p/4 subtraction protocol.

### Data analysis

Conductance, G, was calculated from I-V relations assuming ohmic instantaneous currents, according to:

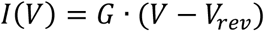

The normalized conductance-voltage (G-V) relations were fit to a Boltzmann function according to equation 1:

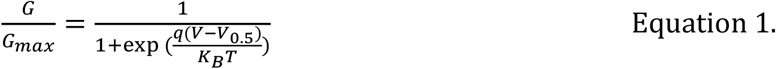

Here, *V*_*0*.*5*_ is the voltage at which *G/G*_*max*_ = 0.5, *q* is the apparent gating charge (in elementary charges, *e*_*o*_) and *K*_*B*_ is the Boltzmann constant and *T* temperature in Kelvin (22°C). The time constant of activation was estimated via a fit of the second half of currents to the equation:

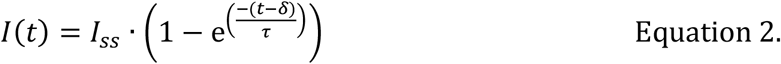

Where *I*_*ss*_ is the amplitude of the current at steady-state, *δ* is the delay of the exponential with respect to the start of the voltage pulse and *τ* is the time constant, both with units of ms. The voltage-dependence of *δ* and *τ* were estimated from a fit to equation:

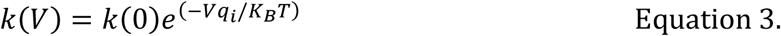

Where *i* stands for *δ or τ* and *k(0)* is the value of either parameter at 0 mV.

Currents in the presence of zinc were normalized to the current before application of the ion to obtain a normalized fraction of current blocked as: F_B_= 1-I/I_max_. The zinc dose response curve was fitted to Hill’s equation in the form:

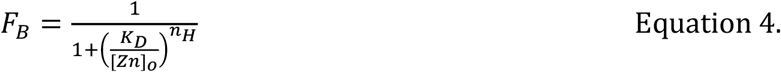

K_D_ is the apparent dissociation constant, [Zn^2+^]_o_ is the extracellular zinc concentration and n_H_ is the Hill coefficient.

## Results

Ion channels have not been studied in corals. In an effort to initiate their study in these organisms, we searched the transcriptome of the Indo-Pacific coral *Acropora millepora* (Moya et al., 2012) for sequences coding for putative voltage-sensing residues present in canonical H_v_1 channels with the form: RxxRxxRIx, which corresponds to the S4 segment of H_v_1 channels and is also found in other voltage-sensitive membrane proteins. Blast searches detected four sequences that seem to correspond to a gene encoding the H_v_1 voltage-activated proton-selective ion channel (Ramsey et al., 2006; Sasaki et al., 2006). *Acropora millepora* is one of the most widely studied species of scleractinian corals and is well represented in the commercial coral trade (Cleves et al., 2018; Ying et al., 2019). We proceeded to clone this gene from a small specimen of *A. millepora* obtained from a local aquarium (Reef Services, Mexico City). As indicated in the methods section, total RNA was extracted from tissue and mRNA was retrotranscribed to obtain cDNA. We managed to obtain a full-length clone and refer to this sequence as AmH_v_1 or H_v_1-type proton channel of *Acropora millepora*.

We were interested in knowing if the same gene is present in a closely related species from the Caribbean Sea. Thus, we used the same primers to clone the H_v_1 channel from *Acropora palmata*, a widespread coral in the same family and which we call ApH_v_1. The amino acid sequence is almost identical to AmH_v_1 (Figure 1-Supplement1A), the greatest divergence is found between a few amino acid residues in the C-terminal region. This result suggests that despite the large biogeographic difference, these two genes have not diverged significantly. The ApH_v_1 sequence also gives rise to fast-activating voltage-gated proton currents (Figure 1-Supplement1B).

The most diagnostic feature of the H_v_1protein is the sequence of the fourth transmembrane domain or S4, which contains three charged amino acids in a characteristic triplet repeat. The presence of these repeats in our sequence allowed us to initially identify our clone as an H_v_1 channel. However, we decided to compare our sequence to those of several H_v_1ortologues. We selected a list of 130 H_v_1protein sequences that are well curated in the Gene Bank (https://www.ncbi.nlm.nih.gov/), representing several branches of the eukaryotes, from unicellular plants to mammals. As expected, the protein sequence of AmH_v_1 has similarity to several other H_v_1 genes from varied organisms (Figure 1A). The identity varies from 98% when compared to other putative coral and anemone sequences, to less than 30% when compared to plant and nematode sequences. In spite of this variability, the putative transmembrane domains of all these proteins show high conservation and consensus sequences logos can detect the presence of highly conserved individual amino acid sequences that can be considered characteristic of H_v_1 channels. Figure 1B compares these transmembrane domain consensus logos with our AmH_v_1 sequence. It can be gleaned that AmH_v_1 contains the highly conserved residues that form the voltage-sensing amino acid residues in S4 as well as their acidic pairs present in S2 and S3. The extracellular histidine residues involved in Zn^2+^ coordination are also present. These results suggest that our sequence is that of a *bona fide* Hv1 voltage-sensing domain (VSD).

**Figure 1.**
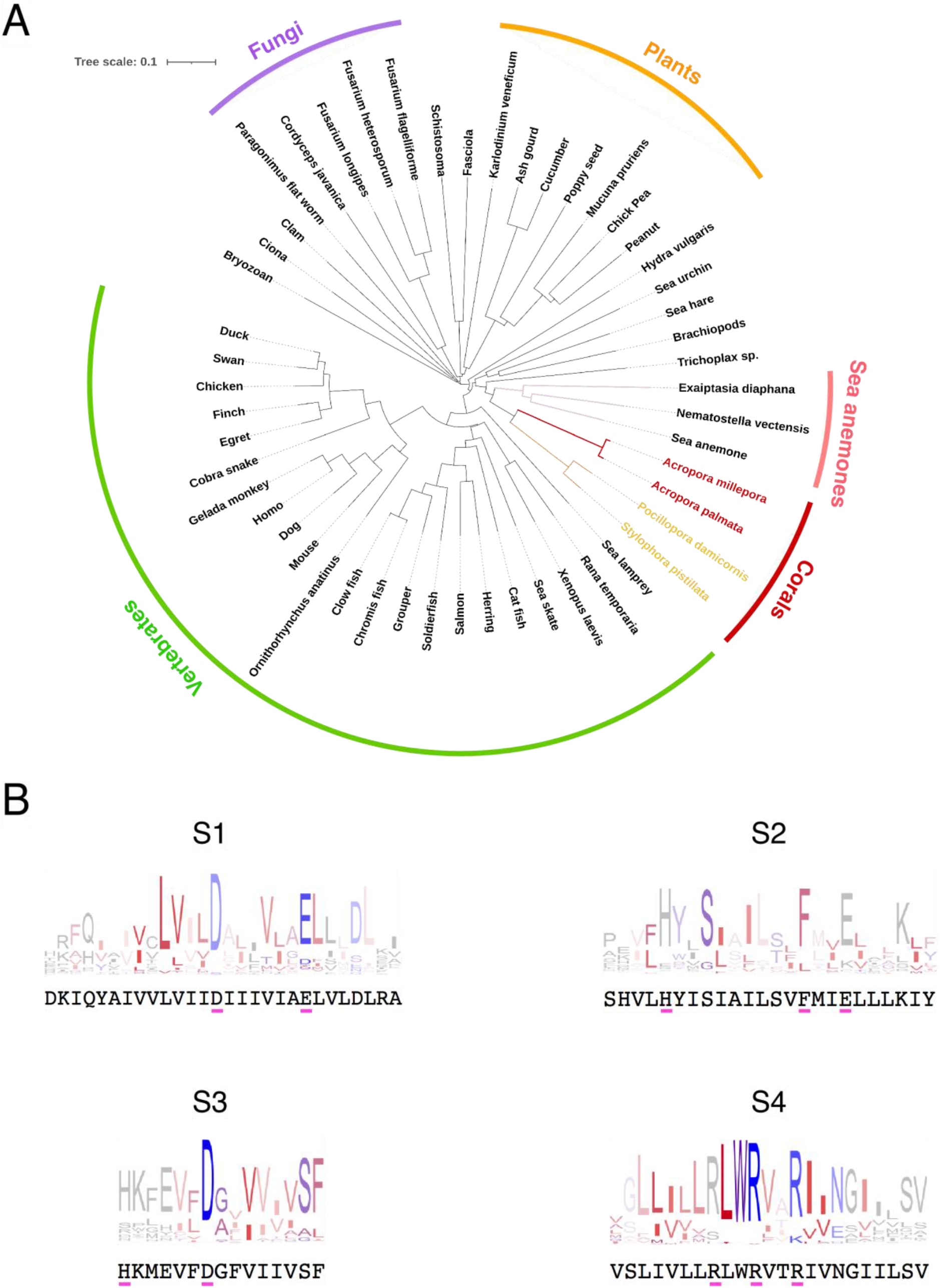
Conservation and phylogenetic relationships of H_v_1 channels. A) Tree obtained from a multiple sequence alignment from H_v_1 channels in CLUSTAL-O. Highlighted in red and yellow are the branches containing coral H_v_1 sequences. B) Consensus logo sequences of transmembrane domains of H_v_1 channels. The color code indicates the hydrophobicity of each residue, where blue indicates charged residues, red indicates non-polar residues and other colors indicate either non-polar or charged residues with less conservation.

Apart from canonical voltage-gated channels, several other proteins contain VSDs. Examples are the voltage-sensing phosphatases like VSPs (Iwasaki et al., 2008) and TPTE and TPTE2 (Halaszovich et al., 2012) proteins (transmembrane protein with Tensin homology) and genes like TMEM266. These proteins are relevant to us since some TPTEs have been shown to also mediate proton currents and TMEM266 can be modulated by Zn^2+^ (Papp et al., 2019). We compared the sequence of AmH_v_1 with several orthologues of TPTEs and TMEM266. Although there is some similarity within transmembrane domains (Figure 1-Supplement 2), the overall sequence comparison shows that AmH_v_1 and these VSD-containing proteins are different.

As mentioned before, we performed a multiple sequence alignment with 130 H_v_1 sequences. In Figure 2 we show the detailed sequence alignment of AmH_v_1 with five of these sequences, which represent some of the best studied H_v_1 genes. It can be seen that there is a high degree of identity, especially in the transmembrane domains. The least degree of conservation appears when comparing this sequence to the dinoflagellate *Karlodinium veneficum* H_v_1 channel (Figure 2A). A search of available transcriptomes from several coral species allowed us to detect the presence of sequences that are found in H_v_1 channels. This suggests that H_v_1 proton channels might be found in many families of scleractinian corals (Figure 2-Supplement 1), as has also been recently shown (Capasso et al., 2021).

**Figure 2.**
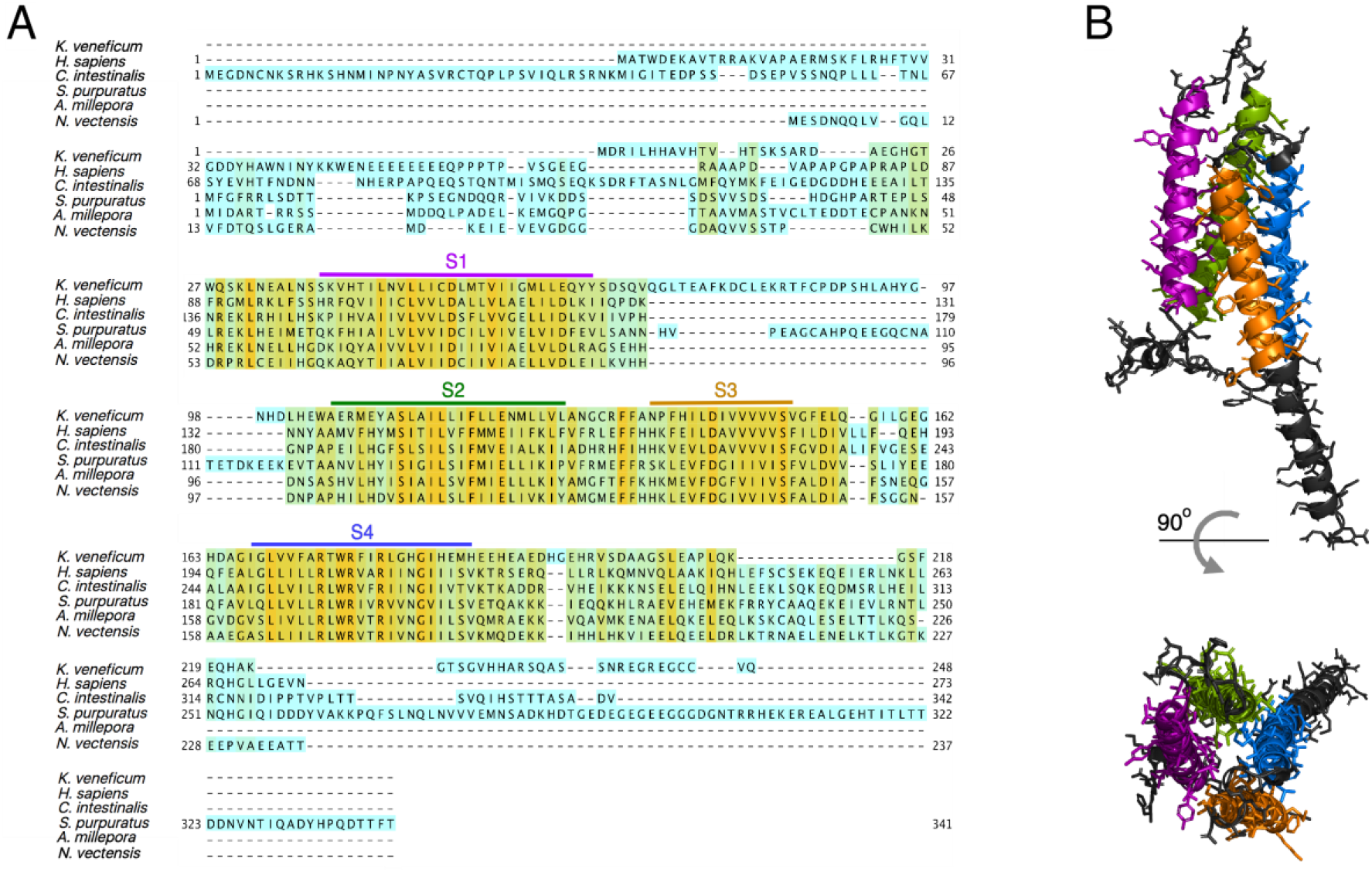
Protein sequence alignment of the *Acropora millepora* H_v_1 (AmH_v_1) channel with selected H_v_1s from other organisms. A) Amino acid sequence alignment of AmH_v_1 with other known H_v_1 orthologues provided by the CLUSTAL-O algorithm. The predicted transmembrane domains are shown by the colored horizontal lines and letters. The colors highlighting the sequence indicate sequence identity. Orange indicates identical amino acids, blue indicates no identity. B) Predicted structural topology of AmH_v_1. Transmembrane domains are colored to correspond with the sequences in A. The top panel is the view parallel to the membrane while the bottom panel is the view from the top (extracellular) side.

Secondary-structure prediction suggests that AmH_v_1 is a canonical H_v_1 channel formed by a voltage-sensing domain with four transmembrane segments. The protein sequence was used for 3D modeling using the SWISS MODEL server (Waterhouse et al., 2018), which produced models based on the H_v_1 chimera structure (Takeshita et al., 2014a) and the Kv1.2 potassium channel voltage-sensing domain (Long et al., 2005). This structural model is shown in Figure 2B. The predicted model indicates a shortened N-terminal region, four transmembrane domains and a long C-terminal helix.

Voltage-gated proton channels from *Ciona* (Sasaki et al., 2006) and humans (Lee et al., 2008a) have been shown to express as dimers in the plasma membrane and this dimeric form is understood to be the functional unit of these proton channels. The dimer is stabilized by a coiled-coil interaction mediated by an alpha helical C-terminal domain. As shown by the model in Figure 2, AmH_v_1 has a long C-terminal helix, which is predicted to engage in a coiled-coil (Paircoil2 (McDonnell et al., 2006). We calculated the probability per residue to form a coiled-coil for all the C-terminal residues, both for human and AmH_v_1 channels, using the program COILS (Lupas et al., 1991). Figure 3A shows that the coiled-coil probability for AmH_v_1 C-terminus is at least as high or higher than that for hH_v_1, an established dimer, strongly suggesting that coral H_v_1s might also form dimers.

**Figure 3.**
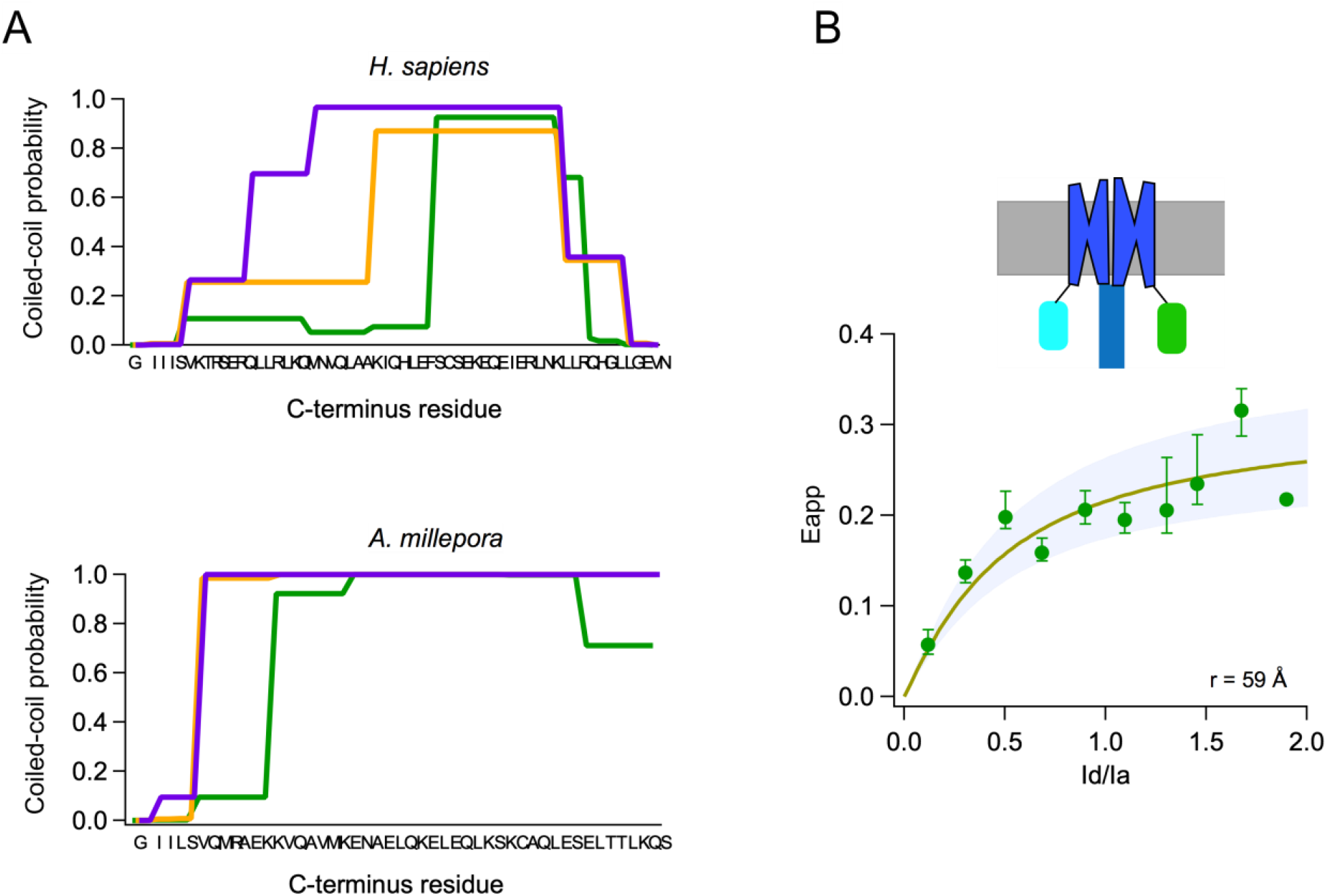
Subunits of the *Acropora millepora* H_v_1 (AmH_v_1) channel associate to form dimers. A) Probability of coiled-coil formation per amino acid residue of the C-terminus domain of hH_v_1 (top) and AmH_v_1 (bottom). The different colors correspond to the three seven-residue windows used by the program to calculate the score. The sequence of the C-terminus is shown in the x-axis. B) FRET measurement of dimer formation. The apparent FRET measured from 134 cells is plotted as a function of the ratio of donor to acceptor fluorescence (I_d_/I_a_). Shown is the average and s.e.m. for data in I_d_/I_a_ windows of 0.1. The continuous curve is the fit of the data to the prediction of a model that considers random assembly of donor- and acceptor-tagged subunits into a dimer. The separation of the FRET pair in a dimer is ∼60 Å, according to the model. The upper panel depicts a cartoon of the presumed FP-tagged dimer in the membrane.

In order to study the oligomeric state of the coral H_v_1, we performed FRET experiments with AmH_v_1 channel tagged with fluorescent proteins as a FRET pair. Figure 3B shows that there is significant FRET efficiency between fluorescent protein-tagged subunits, indicating a very close interaction between monomers. The measured apparent FRET efficiency vs. the fluorescence intensity ratio can be fitted to a model were the subunits assemble as a dimer. From this fit, we can estimate a distance between fluorophores of ∼60 Å, which is compatible with AmH_v_1 being a dimer, at least in HEK293 cells.

### Functional expression of AmH_v_1. Voltage-dependence and kinetics

The cDNA of AmH_v_1 was cloned in the pcDNA3 expression vector and transfected into HEK293 cells. Under whole-cell conditions we recorded large voltage-dependent outward currents. Figure 4A shows a family of such currents. The data suggest that these currents were carried mostly by protons, since the reversal potential, measured from a tail current protocol, closely followed the equilibrium potential for protons, as given by the Nernst equation (Figure 4B).

**Figure 4.**
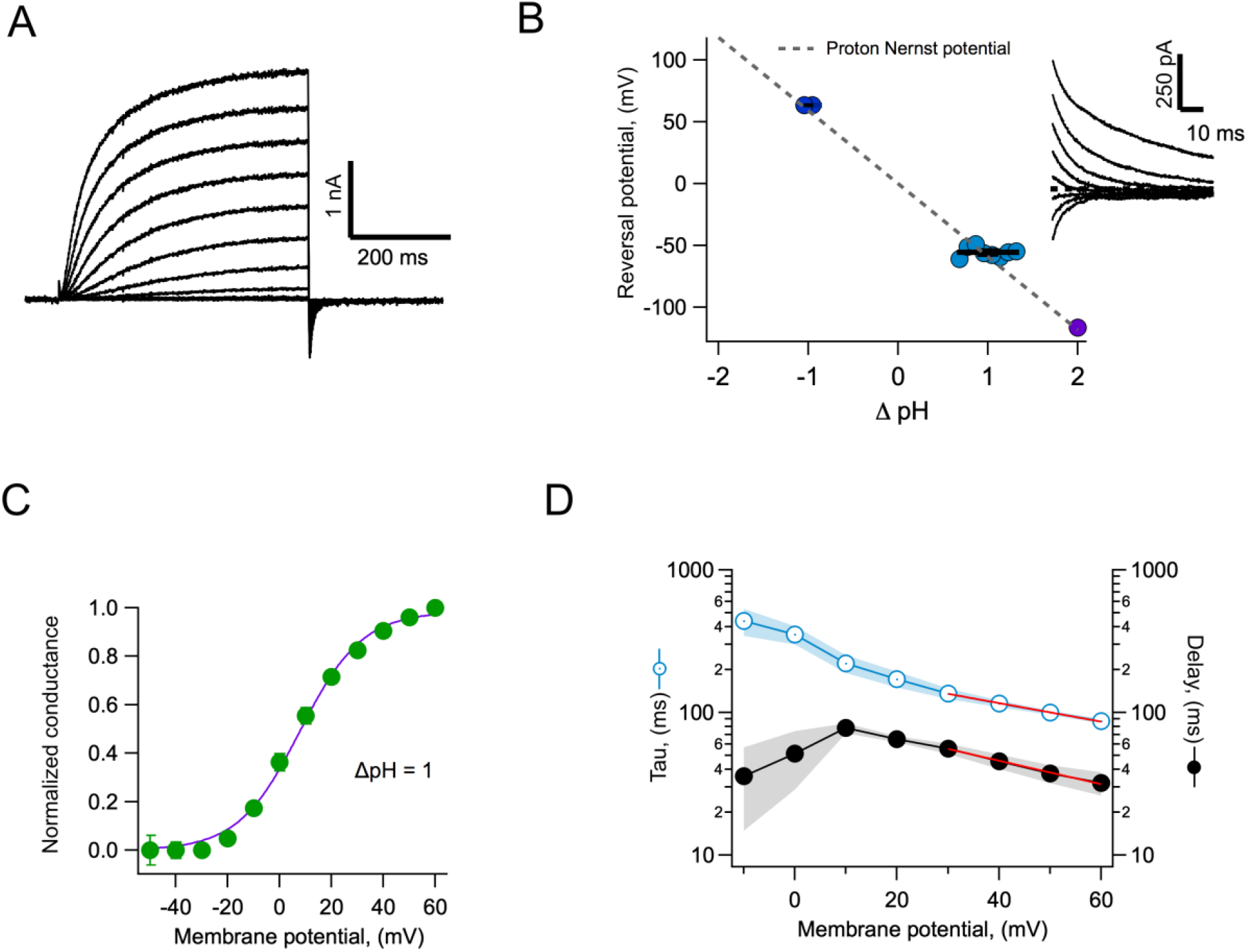
Proton currents mediated by AmH_v_1 expressed in HEK 293 cells. A) Typical proton current family elicited by depolarizing pulses from −50 to 60 mV in 10 mV intervals. The duration of the pulses is 500 ms. Linear current components have been subtracted. B) Reversal potential of currents as a function of the pH gradient. Symbols are individual data and the black horizontal lines are the mean. The dotted line is the expected reversal potential as predicted by the Nernst equation. The inset shows a tail current family from which instantaneous IV curves where extracted to measure the reversal potential. Recordings shown in A and B were obtained in the whole-cell configuration. C) Normalized conductance-voltage curve at ΔpH = 1. The red curve is the fit to equation 1 with parameters V_0.5_ = 7.85 mV, q = 2.09 e_o_. Circles are the mean and error bars are the s.e.m. (n = 7). D) Kinetic parameters of activation. Activation time constant and delay estimated from fits of current traces to equation 2. Circles are the mean and the s.e.m. is indicated by the shaded areas (n = 6). The voltage-dependence of the delay and tau of activation were estimated from a fit to equation 3, which appears as the red curve. Parameters are: δ(0) = 98.2 ms, qδ = 0.47 e_o_. The voltage-dependence parameters for tau are: τ(0) = 212 ms, qτ = 0.37 e_o_.

The voltage-dependence of channel gating was estimated from a fit of the normalized conductance vs. voltage to equation 1. The steepness of the curve corresponds to an apparent charge of ∼2 e_o_, comparable to other H_v_1’s under similar recording conditions (Figure 4C). Interestingly, these channels seem to activate rapidly. This is apparent from the current traces, which reach a steady-state within a few hundred ms (Figure 4A), as quantified in Figure 4D. Equation 3 estimates two parameters, an activation time constant (τ) and a delay (δ). Both the time constant and the delay are similarly voltage-dependent at positive potentials. The existence of a delay in the time course implies that activation is a multiple state process. The delay magnitude is smaller than the time constant at all voltages, which can be interpreted to mean that the rate limiting step for opening comes late in the activation pathway (Schoppa and Sigworth, 1998).

### Comparison to human H_v_1 channel properties

Human H_v_1 is probably the best characterized of the voltage-gated proton channels (Musset et al., 2008), so we compared some of the properties of AmH_v_1 with hH_v_1. AmH_v_1 channels activate faster than their human counterpart. Figure 5 compares the activation kinetics of these two channels under the same conditions. Steady-state is apparently reached sooner after a voltage pulse in AmH_v_1 (Figure 5A) when compared to hH_v_1 (Figure 5B). The slower kinetics of the human orthologue is also evidenced in the more sluggish deactivation tail currents (Figure 5B). The range of voltages over which activation happens is also different between the two channels, with the coral H_v_1 channel activating 40 mV more negative than the human clone (Figure 5C. Notice that the proton gradient is the same in these recordings). Even though AmH_v_1 activates at more negative voltages, the activation range is still more positive than the proton reversal potential, thus coral proton currents activated by depolarization, in the steady state and at least as expressed in HEK293 cells, are always outward.

**Figure 5.**
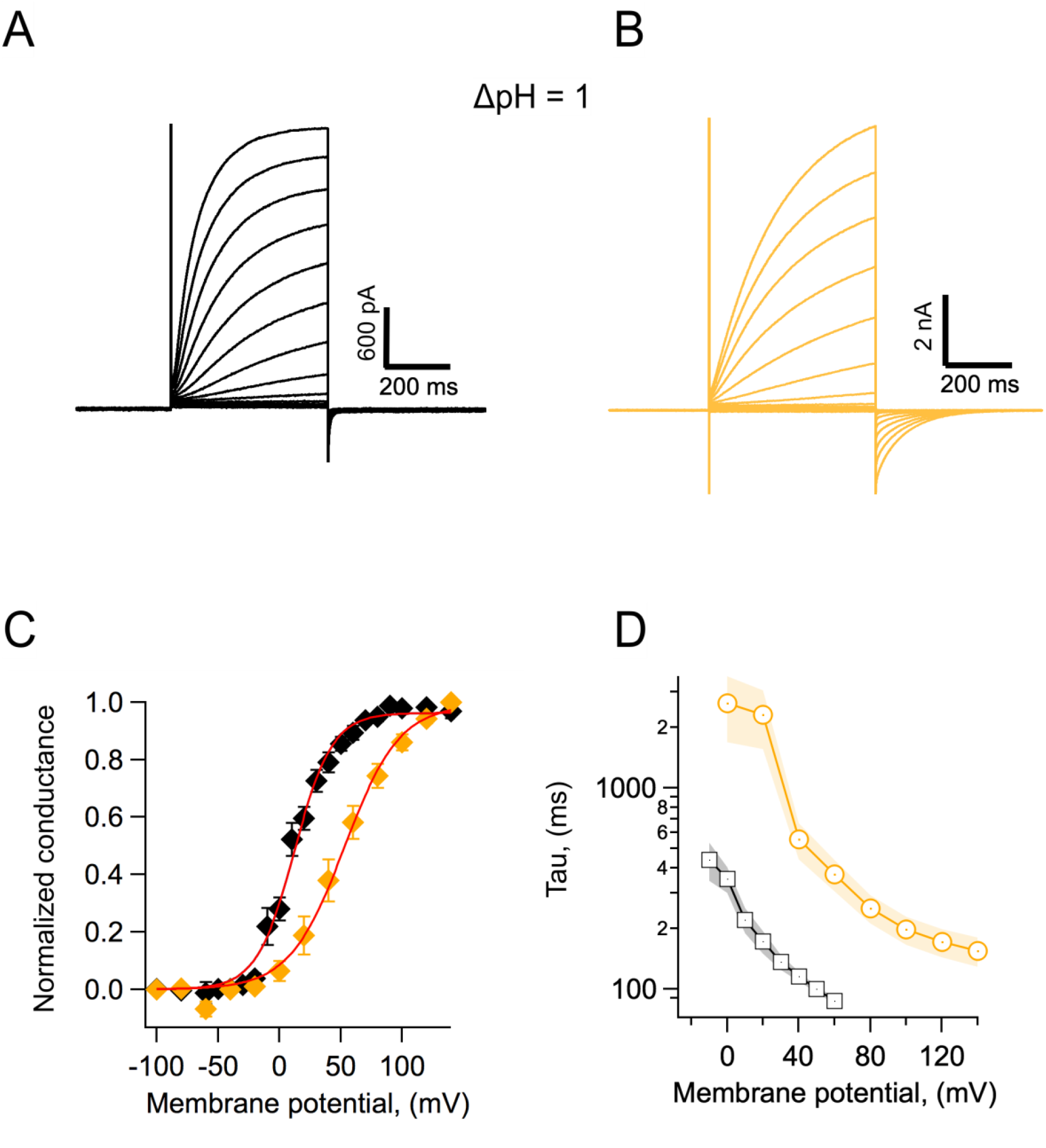
Coral H_v_1 channels are faster and activate more readily than their human counterpart. A) AmH_v_1 currents in response to voltage-clamp pulses from −100 to 120 mV. B) Currents through hH_v_1 channels in response to voltage-clamp pulses from −100 to 120 mV. Recordings shown in A and B were obtained in whole-cell the configuration. C) Comparison of the conductance-voltage relationship for both channels. Black diamonds are the mean G/G_max_ for AmH_v_1 and yellow diamonds for hH_v_1. The error bars are the s.e.m. (n= 3, for both channels). The continuous red curves are fits to equation 1. The fitted parameters are: AmH_v_1, q = 1.62 e_o_, V_0.5_ =12.2 mV; hH_v_1, q = 1.11 e_o_, V_0.5_ = 53.1 mV. D) The activation time constant estimated from fits of currents to equation 2. Circles are the mean for hH_v_1 and squares for AmH_v_1. The shaded areas are the s.e.m. (n= 3, for both channels).

The faster kinetics of AmH_v_1 is clearly evidenced when the time constant of activation, *τ*, estimated using fits of the activation time course to equation 2, is compared for coral and human H_v_1 channels. AmH_v_1 is almost 10-fold faster at 0 mV and over a range of positive voltages (Figure 5D).

### Effects of the pH gradient on gating

Both native and cloned voltage-gated proton channels are characteristically modulated by the pH gradient (Cherny et al., 1995; Sasaki et al., 2006; Ramsey et al., 2006). We carried out experiments to investigate the modulation of the coral H_v_1 channels by different pH gradients. We first recorded whole-cell currents at various ΔpH and estimated the voltage-dependence of the conductance. These G-V curves were fitted to equation 2 to obtain the voltage of half activation, V_0.5_ and apparent gating charge, *q*, that determines the steepness of the fit. As is the case with other H_v_1 channels, the V_0.5_ shifts to negative voltages when ΔpH is greater that 0 and to positive voltages when ΔpH<0 (Figure 6A). When we plot the V_0.5_ as a function of ΔpH the relationship seems to be mostly linear over the range of ΔpH −1 to 2. This relationship is somewhat steeper than the generally observed −40 mV/ΔpH (Figure 6B). We tried to obtain recordings over an extended range of ΔpH values. To this end, we performed inside-out recordings in which the composition of solutions can be better controlled, tend to be more stable and the size of currents is smaller. However, recordings were unstable at extreme pH values and we only managed to reliably extend the data to ΔpH of −2. Figure 6C shows the summary of the inside-out recordings. We have plotted both the V_0.5_ and the threshold voltage, V_Thr_. To obtain this last parameter, we fitted the exponential rise of the G-V curve to a function of the form:

**Figure 6.**
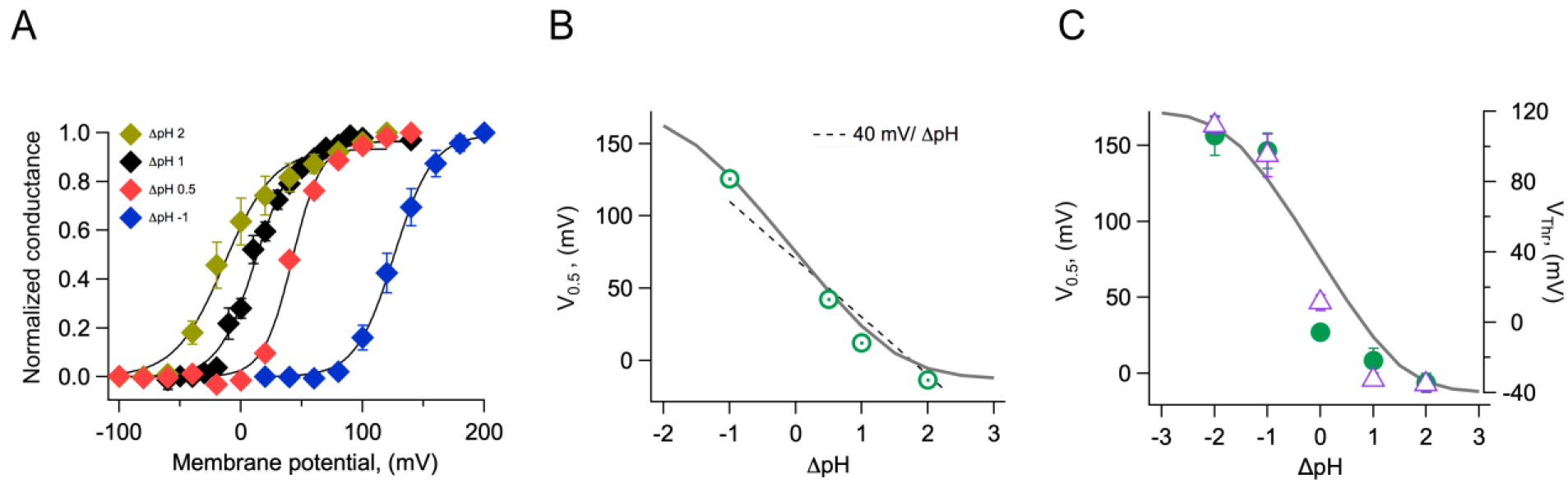
Modulation of channel activation by the pH gradient. A) Conductance vs. voltage relationships obtained at the indicated ΔpH values, obtained from whole-cell recordings of AmH_v_1 proton currents. Continuous lines are fits to equation 1. B) The parameter V_0.5_ was obtained from the fits in A and is displayed as a function of ΔpH. The dotted line is the 40 mV/ΔpH linear relationship. The continuous grey curve is the prediction of the allosteric model. C) Parameters V_0.5_ (green circles) and V_Thr_ (purple triangles) obtained from a different set of inside-out current recordings. Data are mean ± s.e.m. The continuous grey curve is the same prediction of the allosteric model shown in B. The model parameters used to generate the theoretical curve are: E=5×10^5^, D=10^5^, C=0.0002, Kv(0)=0.00005, q_g_=1.0 e_o_, pK_o_=3.4, pK_i_=7.

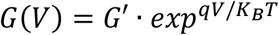

V_Thr_ was calculated as the voltage at which the fit reaches 10 % of the maximum conductance. The parameter V_Thr_ should be less sensitive than V_0.5_ to the possible change in the proton gradient that can occur with large currents. It is clear from these data that at extreme values the dependence of V_0.5_ or V_Thr_ on ΔpH deviates from a simple linear relationship and instead it appears to saturate with increasing ΔpH.

### Allosteric model of voltage and pH-dependent gating

Currently, there are is only one quantitative model that has been used to explain ΔpH gating of H_v_1 channels (Cherny et al., 1995). However, this model is euristic and does not provide mechanistic insight into the process of proton modulation of the voltage dependence of proton permeable channels. In order to explain the modulation of the range of activation by the proton gradient, parameterized by the V_0.5_, we developed a structurally-inspired allosteric model of voltage and proton activation. As many voltage-sensing domains, H_v_1 has two water-occupied cavities exposed to the extracellular and intracellular media (Ramsey et al., 2010; Islas and Sigworth, 2001; Ahern and Horn, 2005). Recent evidence suggests that these cavities function as proton-binding sites through networks of electrostatic interactions (De La Rosa et al., 2018). In our model, we propose that these two proton-binding sites, one intracellular and one extracellular, allosterically modulate the movement of the voltage-sensing S4 segment and thus channel activation in opposite ways. The extracellular site is postulated as inhibitory, while the intracellular site is excitatory, facilitating voltage sensor movement. As a first approximation, we employ a simplified allosteric formalism based on a Monod-Wyman-Changeux (MWC) style model (Horrigan and Aldrich, 2002; Changeux, 2012). As a simplifying assumption, in this model we assume that the voltage sensor moves in a single voltage-dependent activation step. We assume the external and an internal proton-binding sites have simple protonation given by a single pK_a_ value. These sites operate as two allosteric modules and are coupled to the voltage sensor according to coupling factors C and D, respectively. These binding sites in turn interact with each other through the coupling factor E. The modular representations of the model are illustrated in Figure 7A, while the full model depicting all open and closed states with all permissible transitions and the corresponding equilibrium constants for each transition is shown in Figure 7B. Full details of equations derived from these schemes are given in supplementary data.

**Figure 7.**
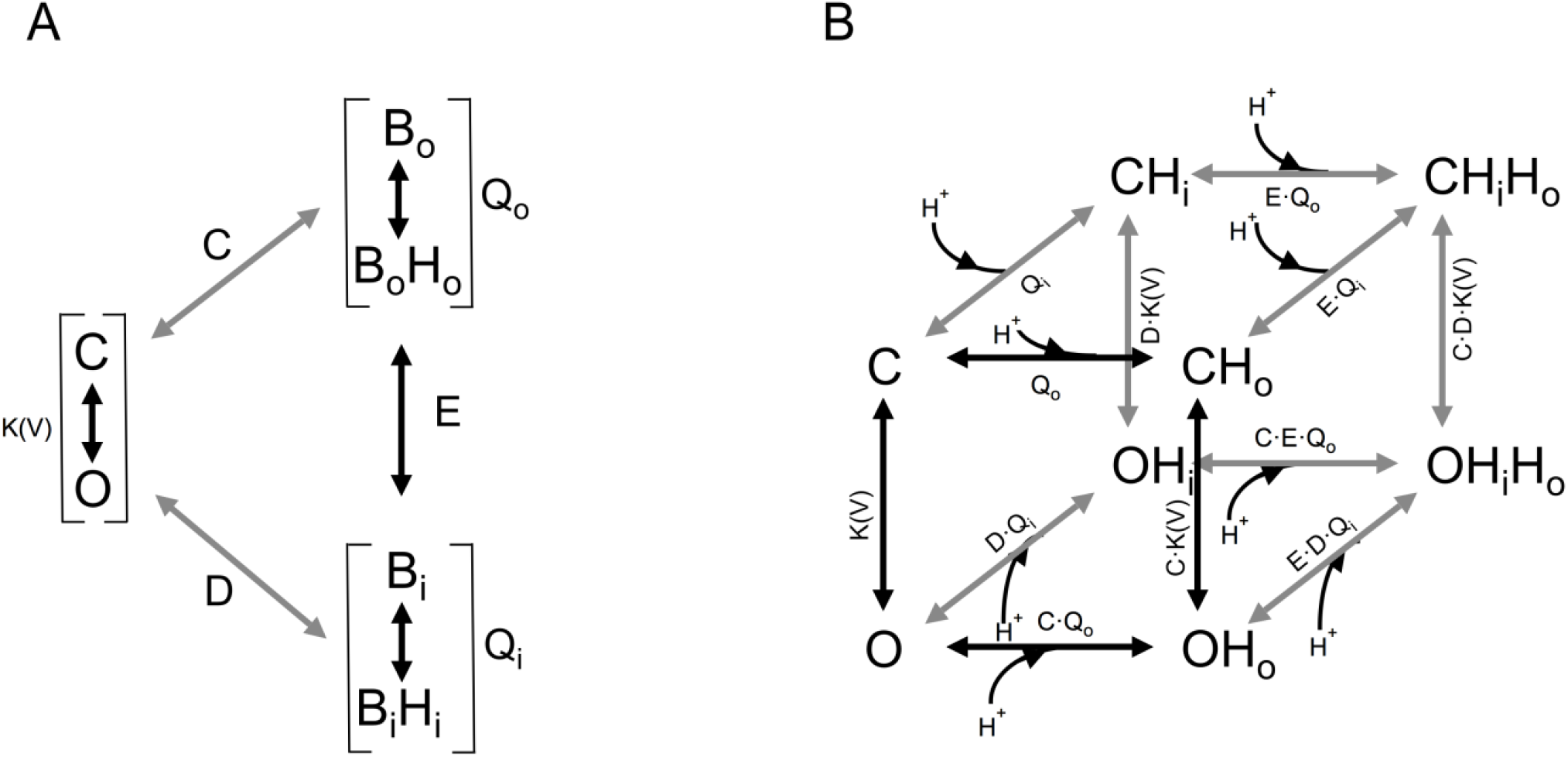
Gating scheme I. A) Modular representation of a simple MWC model; the channel opening transition is voltage-dependent, with equilibrium constant *K(V)*. B_o_ and B_i_ are the unbound states of the extracellular and intracellular proton-binding sites, respectively and B_o_H_o_ and B_i_H_i_ are the proton bound states of these binding sites. Q_o_ and Q_i_ are equilibrium constants that depend on the pK_a_ of each of these binding states. C, D and E are the coupling constants between each of the indicated modules. B) All the individual states implied in A are depicted, along with proton-binding states and the appropriate equilibrium constants. C, closed states, O, open states. OH_x_, OH_x_H_x_ and CH_x_, CH_x_H_x_ are single or doubly proton-occupied states, where x can be o for outside or i for inside-facing binding sites.

This allosteric model represents a first attempt at producing a quantitative mechanistic understanding of the interaction of the voltage sensor and protons in H_v_1 channels.

From the data shown in Figure 6C, it can be seen that the model is capable of reproducing the very steep dependence of V_0.5_ on ΔpH and importantly, the saturation of this relationship at extreme values. Some H_v_1 channels from other organisms show a linear dependence of gating over a large range of ΔpH values, while others show a reduced dependence and even saturation over some range of on ΔpH (Thomas et al., 2018). Our model can explain these different behaviors as different channels having distinct values of pk_a_s for the internal or external sites, differences in coupling factors or differences in the voltage-dependent parameters (Figure6 -Supplement 1).

### Block by Zn^2+^

The best characterized blocker of proton channels is the divalent ion zinc (Cherny et al., 2020; De La Rosa et al., 2018; Qiu et al., 2016). We performed experiments to determine if Am H_v_1 channels are also inhibited by zinc. We found that indeed, extracellular application of zinc in outside-out patches produced inhibition of the channels, reflected in reduced current amplitude (Figure 8A). Figure 8B shows average current-voltage relationships in the absence and presence of 10 μM external zinc. It can be seen that the fraction of current blocked is not the same at every voltage, indicating that this inhibition might be voltage-dependent. The fraction of blocked channels was calculated and is plotted at each voltage along with the I-V curves (Figure 8B). It can be clearly seen that inhibition by Zn^2+^ is voltage-dependent. A simple mechanism for voltage-dependent blockage was proposed by (Woodhull, 1973). This model postulates that a charged blocker molecule interacts with a binding site in the target molecule that is located within the electric field. Fitting the data according to this model, and given that zinc is a divalent ion, its apparent binding site is located at a fraction δ = 0.2 of the membrane electric field from the extracellular side (Figure 8B).

**Figure 8.**
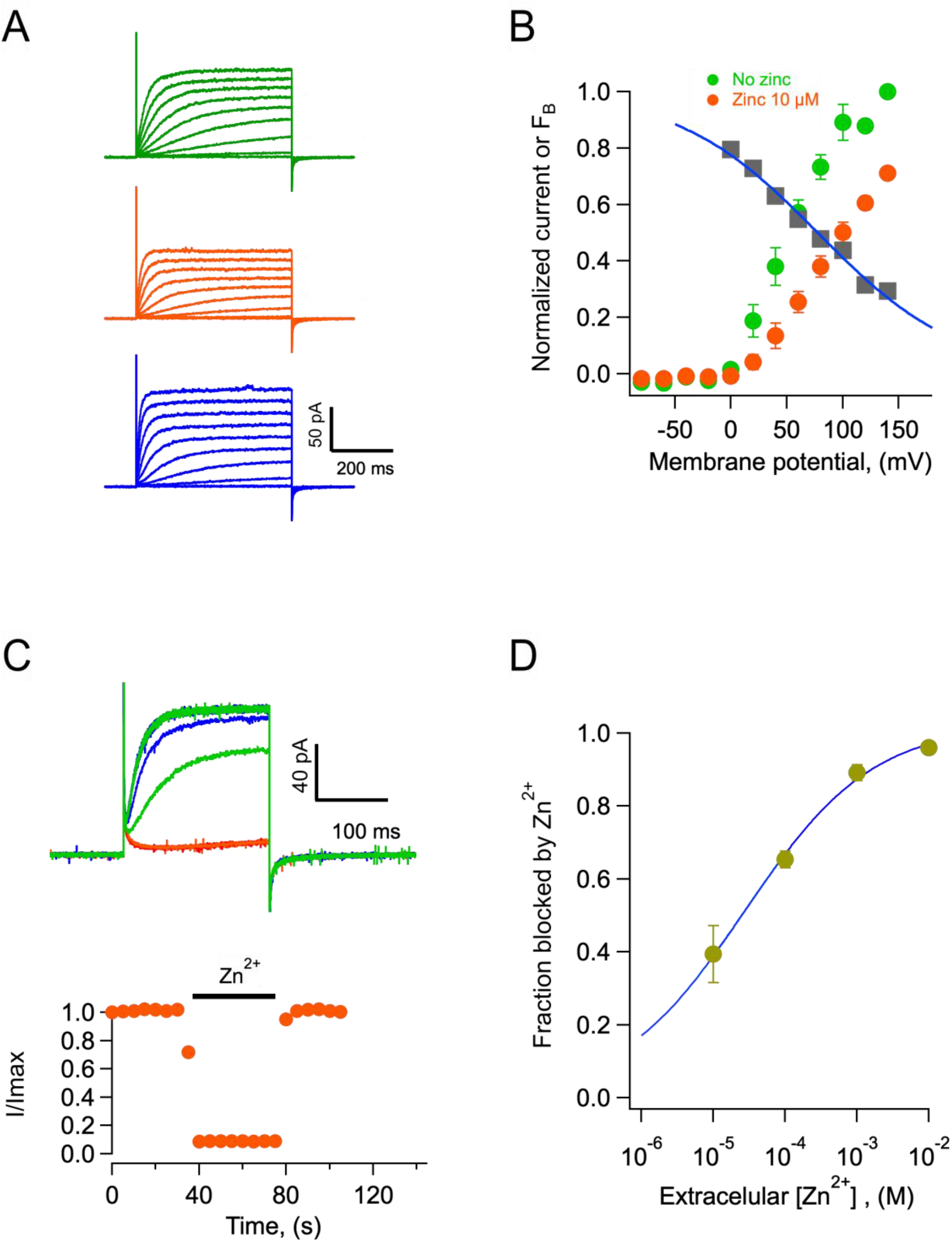
Block of AmH_v_1 channels by extracellular zinc. A) AmH_v_1-mediated currents from an outside-out patch in the absence (top), presence of 10 μM zinc (middle) and after washing of zinc (bottom). The scale bars apply to the three current families. B) Normalized current voltage relationships before and in the presence of 10 μM zinc from 4 patches as in A. The grey squares are the ratio I_zinc_(V)/I(V), which gives the voltage-dependence of the blocking reaction. The blue curve is the fit to the Woodhull equation: 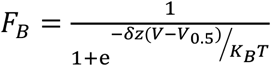, where *F*_*B*_ is the fraction of current blocked, δ is the fraction of the electric field where the blocker binds, *z* is the valence of the blocker, V_0.5_ is the potential where half of the current is blocked, K_B_ is Boltzmann’s constant and T the temperature in Kelvin. The fitting parameters are: δ=0.19, V_0.5_=77.6 mV. C) The effect of zinc is fast. Application of 1 mM zinc to an outside-out patch produces almost instantaneous block of ∼90 % of the current. The effect also washes off quickly upon removal of zinc. Trace colors are as in A. Voltage pulse was 100 mV applied every 5 sec. D) Dose-response curve of zinc block of AmH_v_1 obtained at 100 mV. The continuous curve is a fit of the data to equation 4 with apparent K_D_ = 27.4 μM and n = 0.48.

Zinc blockage proceeds very fast. At 1 mM the channels are blocked almost instantaneously, and the inhibition washes off very fast as well (Figure 8C). Finally, we report the dose response curve (Figure 8D). The inhibition dose response curve can be fit by a Hill equation (Equation 4) with a slope factor of near 0.5 and an apparent dissociation constant, K_D_ of 27 μM.

## Discussion and conclusions

A few ion transport mechanisms in reef-building corals have been described, but up to now, no ion channels have been characterized from any scleractinian species. Here we show that voltage-gated proton-permeable channels formed by the H_v_1 protein are present in corals. In particular, we have cloned these channels from two species of the genus *Acropora, A. millepora and A. palmata*. It is interesting that the protein sequence of these proteins shows a very high degree of conservation, suggesting that, even when the two species are found in different oceans, they haven’t had time to diverge substantially or alternatively, selective pressures on these channels are very similar in both species. The presence of H_v_1 sequences in many other species of corals from disparate clades, suggest that H_v_1 plays an important role in coral physiology.

H_v_1 channels are formed by a protein fold that is structurally equivalent to the voltage-sensing domains (VSDs) of canonical voltage-gated channels (Sasaki et al., 2006; Ramsey et al., 2006). The VSD is formed by a bundle of four antiparallel alpha helices (Takeshita et al., 2014b). In some species, it has been shown that H_v_1 channels are dimeric (Lee et al., 2008b; Mony et al., 2020; Lee et al., 2008a). Accordingly, we also show here that the AmH_v_1 is a dimer. Our FRET results are consistent with the high propensity to form a coiled-coil shown by its C-terminal domain.

H_v_1 channels are different from canonical voltage-gated channels in that both voltage-sensing and permeation are mediated through a single protein domain. Voltage-sensing is thought to occur through the interaction of charged amino acid side chains with the electric field, leading to outward movement of the fourth domain or S4, in a similar fashion to other voltage-sensing domains (Carmona et al., 2018; De La Rosa and Ramsey, 2018). This outward movement of the S4 is coupled to protons moving through the VSD in a manner that is not completely understood (Randolph et al., 2016). Most proton permeable channels seem to have evolved to extrude protons from the cell, and towards this end, their voltage dependence is tightly modulated by the proton gradient between extracellular and intracellular solutions (Cherny et al., 1995).

Our electrophysiology experiments show that these coral channels give rise to proton currents when expressed in HEK293 cells and that they retain the functional characteristics that have been shown to define the class in other species, such as very high selectivity for protons, activation by voltage and modulation of this activation by the proton gradient. The new channels reported here activate faster that the human H_v_1 channel. It has been known that different orthologs of H_v_1 activate with varying kinetics. For example, sea urchin, dinoflagellate and recently, fungal H_v_1 channels activate rapidly, while most mammalian counterparts have slow activation rates (Musset et al., 2008; Smith et al., 2011; Zhao and Tombola, 2021). A comparative study suggests that two amino acids in the S3 transmembrane segment are important determinants of kinetic differences between sea urchin and mouse H_v_1 (Sakata et al., 2016). The authors suggest that the time course of activation is slow in channels containing a histidine and a phenylalanine at positions 164 and 166, respectively (mouse sequence numbering). The AmH_v_1 has a histidine at equivalent position 132 and a methionine at 134. It is possible that this last amino acid in AmH_v_1 confers most of the fast kinetics phenotype. A separate work showed that a lack of the amino-terminal segment in human sperm H_v_1 also produced fast-activating channels (Berger et al., 2017). Interestingly, the *Acropora* channels have a shorter amino-terminal sequence, which could also contribute to their fast kinetics.

One of the most interesting characteristics found in these new proton channels is their modulation by the proton gradient. As opposed to other H_v_1 channels, we can observe a trend towards saturation of the V_0.5_ for activation as a function of ΔpH at extreme values of this variable. A tendency towards saturation of the V_0.5_-ΔpH relationship has been observed in mutants of the hH_v_1 channel (Cherny et al., 2015) or at negative values of ΔpH for a snail H_v_1 (Thomas et al., 2018), but it seems it can be fully appreciated in AmH_v_1. Since our model explains the observation of saturation of voltage gating at extreme values of ΔpH as a consequence of the existence of two saturable sites for proton binding, we attribute this behavior, to the large separation of pK_a_ values for the extracellular and intracellular proton binding sites.

The strength of allosteric coupling of these sites and the voltage sensor will determine if saturation is observed over a short or extended range of ΔpH values and the range of values of V_0.5_ that a particular channel can visit. Our model should provide a framework to better understand gating mechanisms in future work.

It is clear that more complicated models, with a larger number of voltage dependent and independent steps (Villalba-Galea, 2014) and coupling to protonation sites should be the next step to improve data fitting and explore voltage-and proton-dependent kinetics. In particular, these types of models can help explain mutagenesis experiments exploring the nature of the protonation sites.

H_v_1 proton channels seem fundamental in handling fluctuations in intracellular pH and take part in several well-characterized physiological processes that depend on proton concentration changes, such as intracellular pH regulation, sperm flagellum beating, reactive oxygen species production and bacterial killing in immune cells, initiation of bioluminescence in single-celled algae, etc. (Castillo et al., 2015).

What is the function of voltage-gated proton channels in corals? The deposition of a CaCO_3_ exoskeleton is one of the main defining characteristics of scleractinians, however, the ionic transport mechanisms involved in this process are mostly unknown. In order for aragonite precipitation to occur favorably, the pH of the calicoblastic fluid, right next to the skeleton is maintained at high levels, between 8.5 and 9 and above the pH of sea water (Le Goff et al., 2017). It has been posited that corals control this pH via vectorial transport of protons to the gastrodermal cavity (Jokiel, 2013). Since proton transport away from the site of calcification would incur a drastically lower intracellular pH in the cells of the aboral region, we propose that, given their ability to rapidly regulate intracellular pH (De la Rosa et al., 2016), H_v_1 proton channels contribute by transporting protons from the cells. Thus, these proton channels would be a major component of the mechanisms of intracellular pH regulation. Given that the activation range of H_v_1 is controlled by the pH gradient, a large intracellular acidification would facilitate opening of these channels at the resting potential of cells, which is presumably negative.

The finding that coral H_v_1 channels retain their sensitivity to Zn^2+^, opens the possibility of using this ion as a pharmacological tool to study the role of proton channels in pH homeostasis. It is interesting that a recent report has shown detrimental effects of zinc supplementation in coral growth (Tijssen et al., 2017), a result that could be explained by zinc inhibition of H_v_1.

The physiological role of H_v_1 channels in corals might be essential in the response of these organisms to ocean acidification. We theorize that as the pH of sea water acidifies, gating of H_v_1 should require stronger depolarization, thus hindering its capacity to transport protons from the cell. This will contribute to a diminished calcification rate and less aragonite saturation of the CaCO_3_ skeleton. It would be interesting and important to study the effects of acidification on H_v_1 physiology and pH regulation in corals in vivo. Essentially nothing is known about the electrophysiological properties of coral cells. This report represents the first time that an ion channel has been cloned and characterized in any coral and should open a new avenue of research, such as uncovering the cellular and possible subcellular localization of these channels and carefully measuring their physiological role in vivo.

## Acknowledgments

We would like to thank Alejandra Llorente for excellent technical assistance. This work was funded in part by grant No. IN215621 from DGAPA-PAPIIT-UNAM to L.D.I., grant No. 247765 to A.T.B. and grant No. IN200720 to T.R. EM was funded by Conacyt-Fronteras en la Ciencia Grant No. 2.

## Author contributions

G.R-Y, performed cloning, performed heterologous expression, performed experiments, read the paper. C.C., M.A.C.-R., E.S-D., performed cloning, expression and electrophysiology and FRET experiments. L.D.I. obtained funding, conceived research, procured animals, analyzed data, wrote the paper. A.B. and E.M. procured collection permits and specimens, performed RNA extraction, revised and edited the paper. I.S.R. contributed ideas, revised and edited the paper. T.R. analyzed data, wrote and edited the paper, contributed ideas.

## Supplementary materials

**Figure 1 - Supplement 1.**
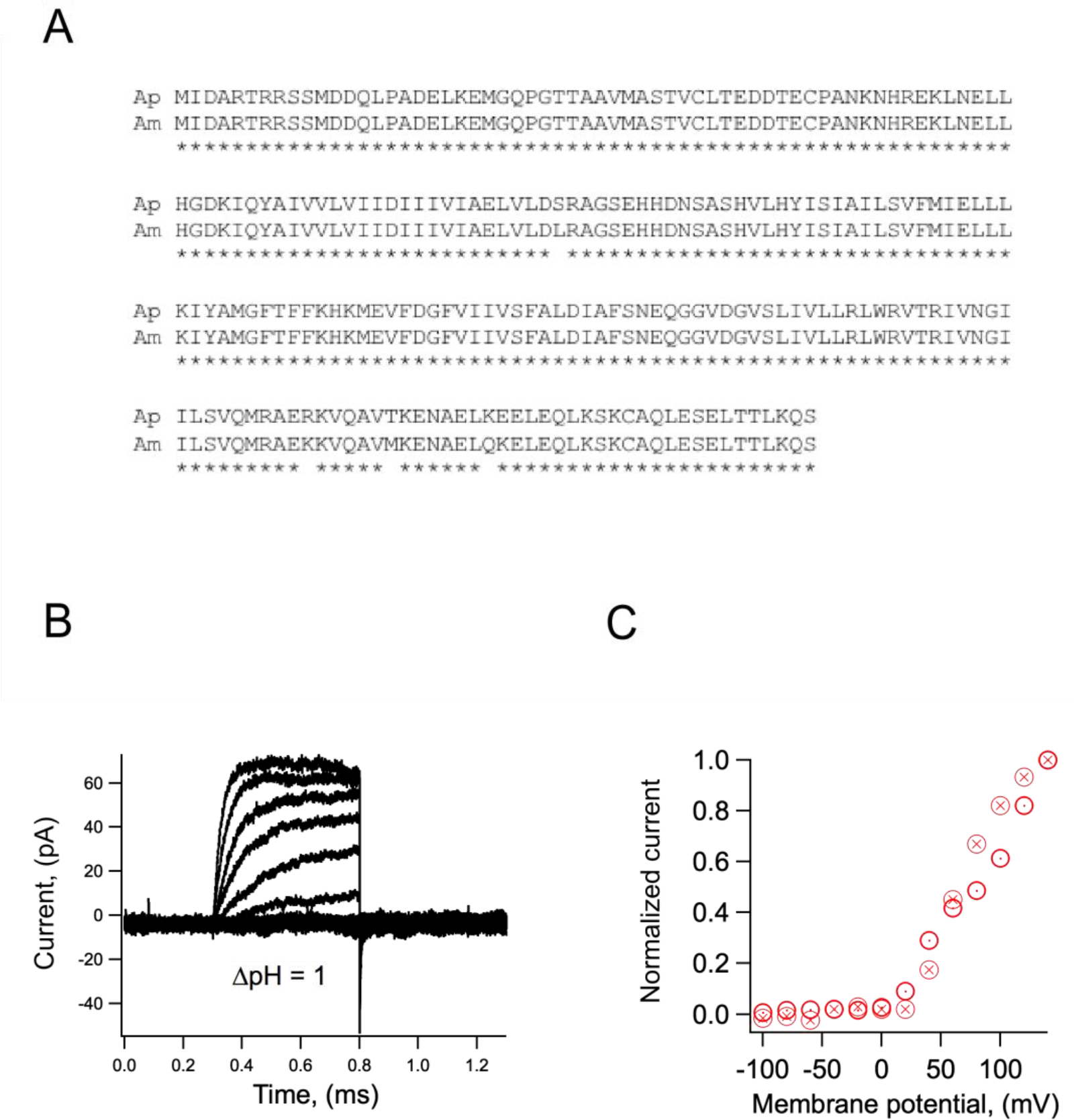
Some characteristics of the H_v_1 from *Acropora palmata*. A) Comparison of the amino acid sequence of H_v_1s from *Acropora millepora* (Am) and *Acropora palmata* (Ap). The asterisks bellow each residue indicate identity. B) Currents elicited from an inside-out patch obtained from a HEK293 cell expressing ApH_v_1. Voltage pulses were from −100 to 140 mV in 20 mV steps. The ΔpH was 1. C) Normalized current-voltage relationships of two patches obtained as in B.

**Figure 1 - Supplement 2.**
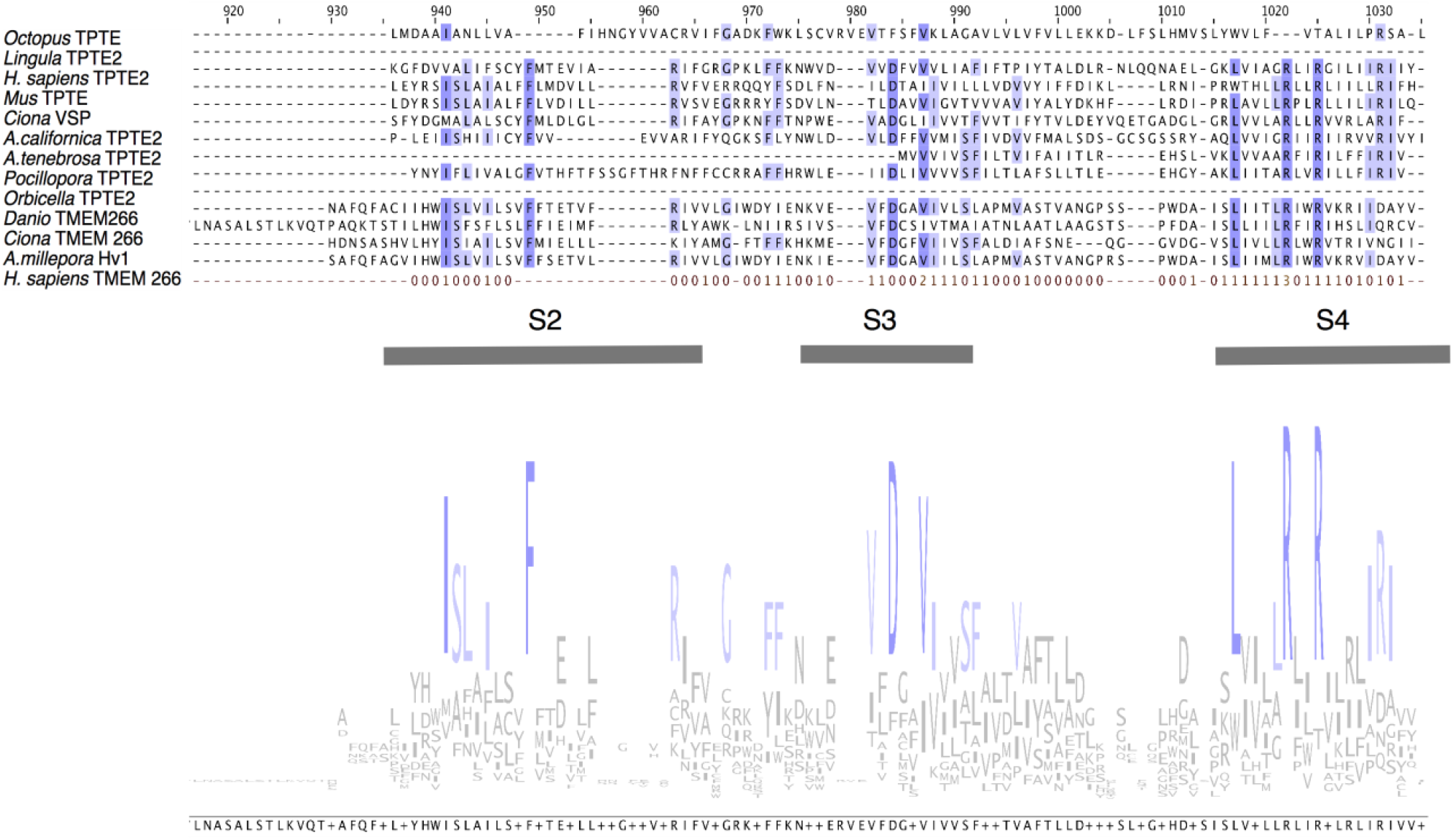
Comparison of the sequence of AmHv1 to other voltage-sensing proteins. Voltage-sensing phosphatases live CiVSP, TPTE and TPTE2 membrane proteins contain a voltage-sensing domain (VSD). Other proteins such as TMEM 266 also contain VSDs. Sequence similarity can be detected in the putative transmembrane domains. We show the sequence alignment of the region with highest similarity between the selected sequences of diverse organisms. The logo consensus sequence shows conservation, specially of the S4 segment. Other amino acid residues common in VSDs are also conserved in S2 and S3. However, TPTE and TMEM266 proteins are very different from *Acropora millepora* H_v_1.

**Figure 2- Supplement 1.**
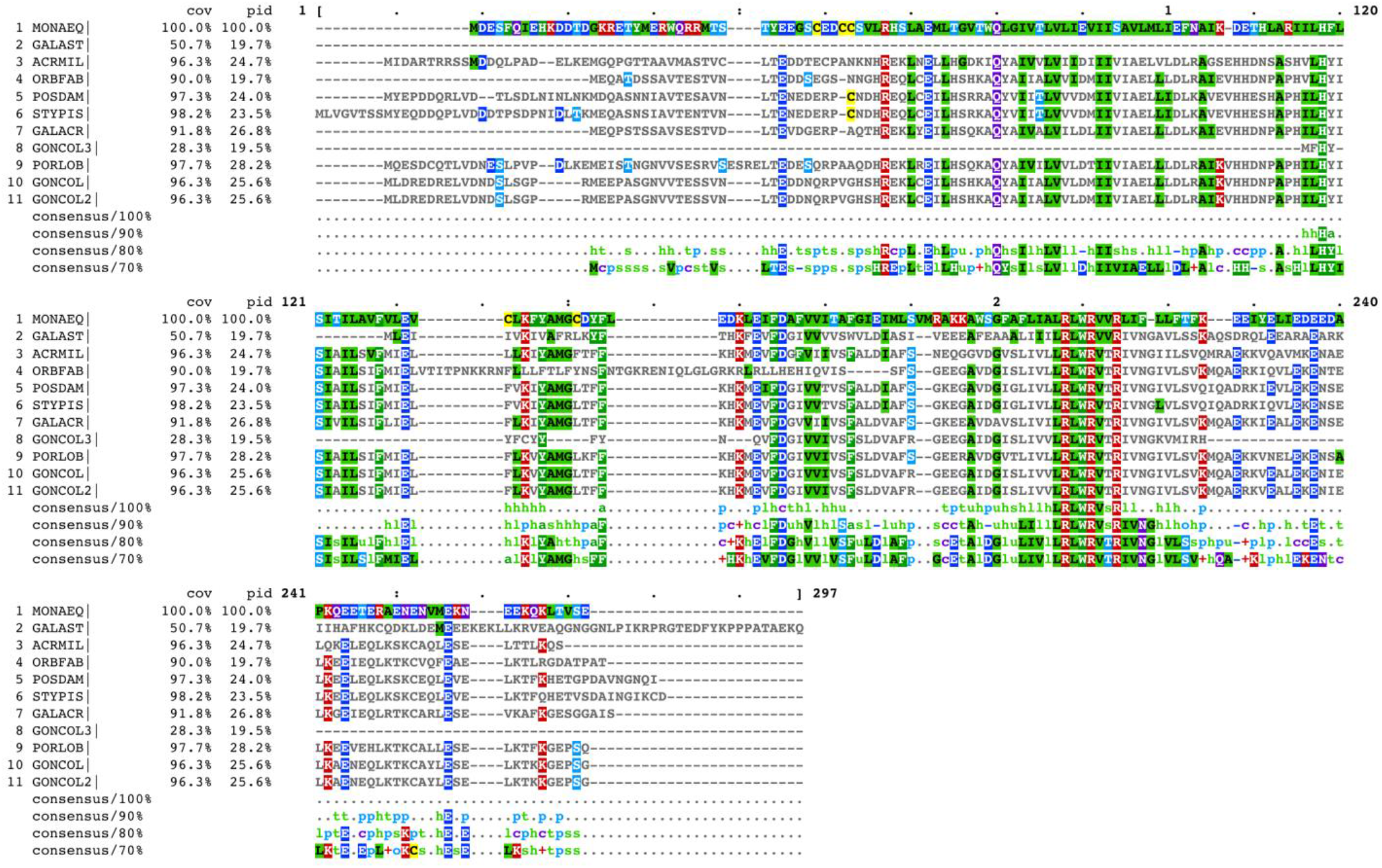
Comparison of the AmH_v_1 protein sequence with similar sequences found in other coral species. MONAEQ: *Montastrea*. GALAST:*Galastrea*. ACRMIL: *Acropora millepora*. ORBFAB: *Orbicela faveolata*. POSDAM: *Posillopora damicornis*. STYPIS: *Stylophora pistilata*. GALACR: *Galaxea*. GONCOL: *Goniopora*. PORLOB: *Porites lobata*.

**Figure 6- Supplement 1. Model equations and simulations**.

The full complement of discrete states in our model is shown in Figure 7B. This allosteric model predicts that the open probability, *P(V, pH)* is dependent on voltage and pH according to the following equations:

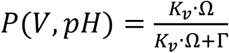

Where:

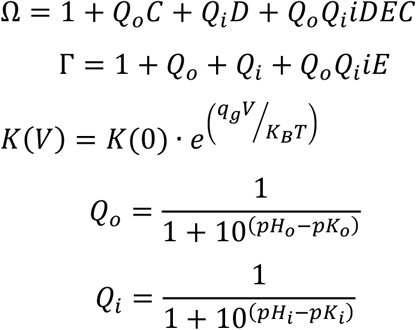

The voltage of half activation is given by:

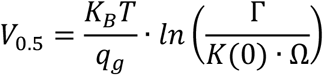

pK_o_ and pK_i_ are the pK_a_ values of the extracellular and intracellular proton binding sites, respectively.

**Supplementary data-Figure 1.**
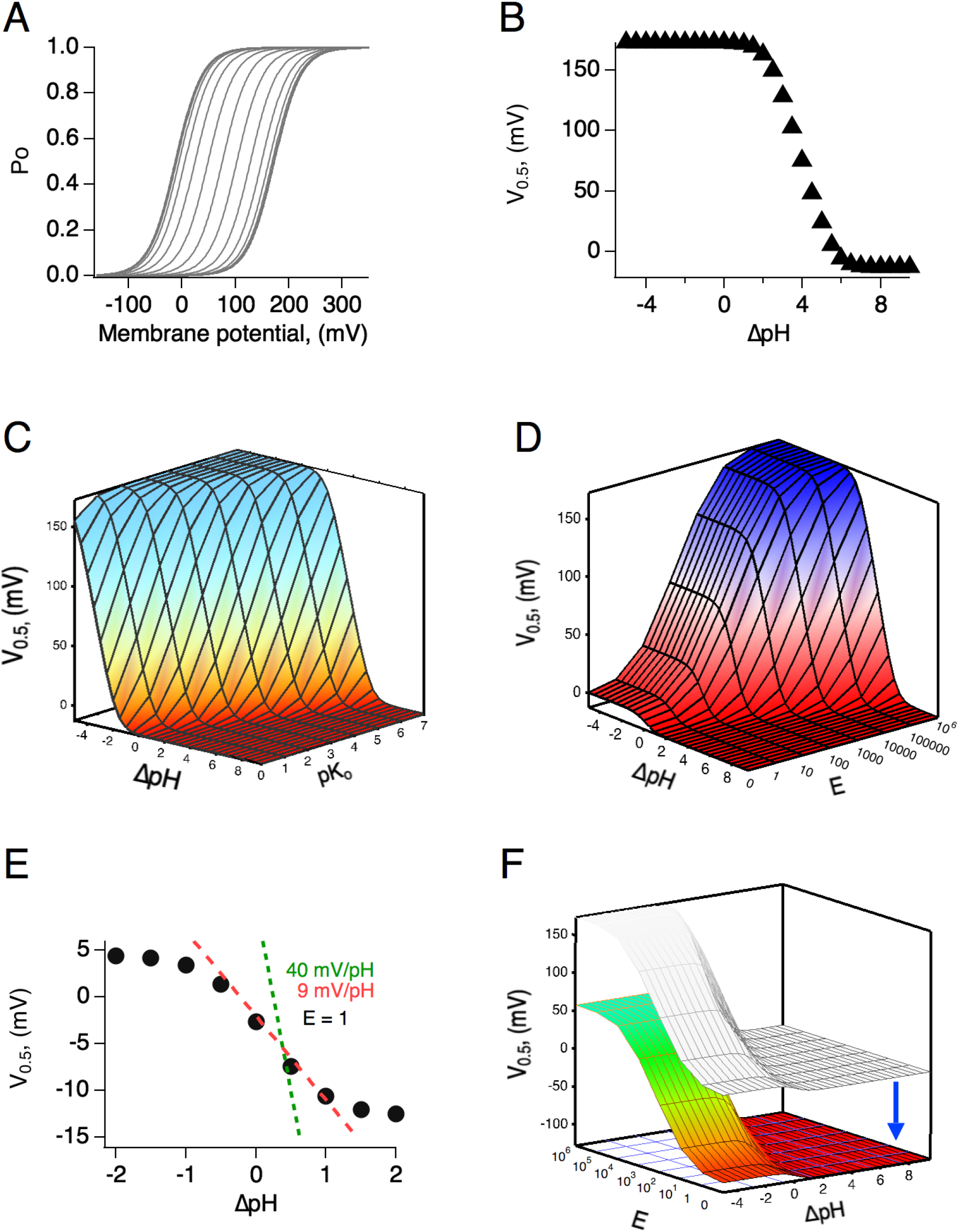
Simulations of the voltage- and pH-dependent behavior predicted by the allosteric model. A) Calculated G-V curves and B) V_0.5_ as a function of ΔpH. Model parameters are: pK_o_, pK_i_ = 7, K(0) = 0.00005, q = 1 e_o_, E = 10^6^, C = 0.0002, D = 10^5^. C) V_0.5_ as a function of ΔpH calculated for different values of the pK_o_. D) V_0.5_ as a function of ΔpH calculated for different values of the coupling factor E, which determines the allosteric communication between external and internal protonation sites. Note that lack of coupling between sites (E = 1), results in channels with very shallow modulation by pH, as illustrated in E. All other parameters are as in A and B. E) A slice of the surface in D, with E =1. The two lines show the expected dependence of V_0.5_ on ΔpH for a fully modulated channel (E>100). Also shown is the dependence of 9 mV/ΔpH unit. F) The range of V_0.5 is_ dependent on the value of the voltage-dependent equilibrium constant at 0 mV. An increase to K(0)=0.005 shifts the whole surface by approximately −110 mV. All other parameters are as in D.

